# Examination of Onset Trajectories and Persistence of Binge-Like Eating Behavior in Mice after Intermittent Palatable Food Exposure

**DOI:** 10.1101/2022.09.28.510003

**Authors:** Britny A. Hildebrandt, Hayley Fisher, Susanne E. Ahmari

## Abstract

Binge eating (BE) is a persistent behavior associated with a chronic course of illness and poor treatment outcomes. While clinical research is unable to capture the full course of BE, pre-clinical approaches offer the opportunity to examine binge-like eating from onset through chronic durations, allowing identification of factors contributing to BE persistence. The current study quantified the trajectories of binge-like eating onset and modeled cycles of abstinence/relapse to develop a translational model for BE persistence. Adult male and female C57Bl6/J mice were randomized to a binge-like palatable food (PF) access schedule (daily 2-hour, 3x/week) or continuous, non-binge like PF access for 12 days (Experiment 1). Persistence of PF consumption in both binge-like PF access groups was then examined across three cycles of forced abstinence and re-exposure to PF (incubation) to model the persistence of BE in clinical populations. Mice with daily 2 hour PF access escalated their intake more than mice in the 3x/week or continuous groups (Experiment 1).This pattern was more pronounced in females. In addition, this pattern of PF intake re-emerged across multiple cycles of behavioral incubation (Experiment 2). These findings provide a model of binge-like eating in mice that can be used in future studies examining both environmental factors and neural mechanisms contributing to BE persistence.

---

Binge eating (BE) is a compulsive behavior with a chronic and persistent course (Pope et al., 2006). Though BE age of onset is often characterized as early adulthood (between 17-22 years of age) (Solmi et al., 2022), a significantly younger onset has also been suggested during childhood and early adolescence (Kjeldbjerg & Clausen, 2021). Critically, longer trajectories of BE illness are associated with poorer treatment outcomes (Vall & Wade, 2015). Despite the significant impact of BE on quality of life and its typical chronicity (Agh et al., 2015), little is known about the mechanisms underlying the persistence of BE. Approaches that model trajectories of BE from onset through chronic persistence are needed to determine these mechanisms.

Current limitations of clinical human research prohibit capturing a complete duration of BE illness from the first episode across the often chronic course of illness (Keski-Rahkonen, 2021). Large cohort studies recruiting individuals before the development of BE are unable to predict how many participants will eventually engage in the behavior; and longitudinal studies recruit individuals already engaging in BE, missing important factors associated with onset. Because of these limitations, it is challenging to isolate neural and behavioral changes associated with the complete trajectory of BE development. However, pre-clinical methods offer a unique opportunity to longitudinally examine binge-like eating from the first episode through chronic duration.

Current pre-clinical approaches for examining binge-like eating in rodents have successfully modeled core components of the clinical presentation of BE (as reviewed in Hildebrandt & Ahmari, 2021). These paradigms generate binge-like eating in rodents through intermittent (rather than continuous) exposure to highly palatable food (PF; i.e., typically high in sweetness, and low in nutritional value) (Boggiano et al., 2007; Furlong et al., 2014; Hildebrandt et al., 2014; Hildebrandt et al., 2018). Additionally, these models yield large amounts of PF consumed within a set short period of time (approximately two hours), modeling the typical phenotype seen in humans (American Psychiatric Association, 2013); and precise quantification of food intake is also possible. This prior work has provided a significant foundation of modeling binge-like eating in rodents that has advanced our understanding of behavioral and neural mechanisms underlying BE episodes.

Despite these advances, no pre-clinical work to date has successfully modeled persistence of binge-like eating over time, limiting translation to humans. Specifically, after childhood onset, BE tends to persist or worsen over time rather than remit/improve (Goldschmidt et al., 2016). Earlier age of BE onset is also associated with poorer treatment outcomes (Safer et al., 2002), leading to a chronic course of illness that persists into adulthood. Additionally, there are moderate rates of relapse (~30%) after cognitive behavioral treatment (Fairburn et al., 1993) and pharmacological treatment (Hudson et al., 2017), suggesting that BE persistence remains an important clinical target even after effective treatment intervention. Pre-clinical models that display not only the onset and development of BE, but also the cyclic nature of remission and re-emergence, are therefore necessary to identify mechanisms contributing to BE persistence that may serve as treatment targets.

The aim of the current study was to develop a pre-clinical model of BE persistence in mice to mimic the cyclical nature of clinical BE. To achieve this goal, we first examined the onset and early intake trajectories of PF in two different binge-like intermittent PF access schedules (daily intermittent, 3x/week) compared to non-binge like continuous PF access in both male and female mice. Second, we examined the persistence of these binge-like PF access schedules across multiple cycles of incubation and re-exposure to PF to mimic the reemergence of BE seen in humans after treatment (Fairburn et al., 1993). We found that daily intermittent PF access, rather than 3x/week or continuous access, leads to the most robust increase in PF intake reflective of BE. Additionally, though persistent patterns of PF intake were observed in animals in both the daily intermittent and 3x/week groups, the daily intermittent group continued to consume the highest amounts of PF during testing. All patterns of binge-like eating were stronger in female versus male mice. Together, these findings help validate a pre-clinical model that can be used as a foundation for future studies targeting the mechanisms underlying BE persistence.

## Materials and Methods

### Animals

Adult male and female C57BL6/J mice (Jackson Laboratories or in-house bred) were used for all experiments. Mice were group housed with 3-5 mice per cage and given ad *libitum* access to standard chow and water for the entirety of the study. Animals were maintained on a 12/12 hour light-dark cycle (lights on at 7:00 AM; off at 7:00 PM). All experiments were approved by the Institutional Animal Care and Use Committee at the University of Pittsburgh in compliance with National Institutes of Health guidelines for the care and use of laboratory animals.

### Experiment 1: Examination of early trajectories of binge-like eating in male and female mice

The BE paradigm used across all experiments is based on previous work using intermittent PF access schedules in rodents (see review in Hildebrandt & Ahmari, 2021). Intermittent PF access schedules mimic multiple features of BE in humans– e.g., BE is episodic and occurs over a short (~two hour) period of time (American Psychiatric Association, 2013), and individuals consume food that is typically low in nutrition but high in palatability (Boggiano et al., 2007; Furlong et al., 2014; Hildebrandt et al., 2014; Hildebrandt et al., 2018). For the first experiment, male and female C57BL6/J mice were randomized to one of three groups: 1) Daily intermittent (N = 3 male, 5 female), in which animals received binge-like intermittent access to PF daily for 2 hours; 2) 3x/week (N = 3 male, 5 female), in which animals received binge-like intermittent access to PF on Monday, Wednesday, and Friday for 2 hours; or 3) Continuous access control (N = 4 male, 4 female), in which animals received ongoing continuous (24 hour), non-intermittent access to PF. The PF used in this paradigm was sweetened condensed milk (Nestle), diluted in a 3:1 ratio with water (Furlong et al., 2014). All mice had *ad libitum* access to standard chow and water throughout the paradigm regardless of access schedule, and animals were never placed on food restriction.

Mice were removed from group-housed conditions, weighed, and placed in individual cages for the duration of each 2-hour feeding test. Feeding tests occurred each day of the paradigm for the continuous and daily intermittent groups, and every Monday/Wednesday/Friday for the 3x/week group. PF was provided in a 50-milliliter conical tube with sipper attachment. Chow was available *ad libitum* for the duration of the feeding test. PF and chow were weighed at the beginning and end of the 2-hour feeding test. Mice were then returned to group home cages. Continuous access mice also had a 50-milliliter tube containing sweetened condensed milk in the home cage for the other 22-hours per day (i.e. when not performing feeding tests), and this tube was weighed daily. This schedule was executed for 12 days resulting in a total of 12 feeding tests for the daily intermittent group, 6 feeding tests for the 3x/week group, and 12 feeding tests plus 12 days of 24-hour PF exposure in the continuous group.

### Experiment 2: Incubation and persistence trajectories of binge-like eating phenotypes

In a new cohort, adult male and female C57BL6/J mice were randomized to one of the two binge-like PF access groups described above: 1) Daily intermittent 2-hour (N = 3 male, 4 female), or 2) 3x/week (N = 3 male, 5 female) to examine persistence of BE phenotypes over time. After completion of a 12 day period of feeding tests following procedures described above in Experiment 1 (Baseline phase), animals remained in group home cages with no feeding tests for 5 days. During this 5 day period without access to PF, animals continued to have access to chow and water. Animals were never placed on food restriction. On Day 6, animals engaged in an additional five days of feeding tests (Incubation Phase), which included a total of five feeding tests for daily intermittent animals and three feeding tests for 3x/week animals. This cycle of forced abstinence and re-exposure to PF was repeated for two more 5-day cycles (total of three 5-day cycles during Incubation Phase). Body weight, PF intake, and chow intake were measured as described above.

### Statistical Analyses

To account for differences in body weight between male and female mice, all food intake (PF and chow) was standardized by body weight prior to analysis (standardized intake = [grams of PF or chow intake] / [body weight in kilograms]). For all BE paradigm data (PF intake, chow intake, body weight), multilevel regressions were run using a full factorial analysis and all categorical predictors were effect coded. For all analyses, the residual distributions were examined for normality and homoscedasticity. All group analysis models met the assumptions. For Experiment 1, predictors included the between-subjects categorical variables Group (continuous, daily intermittent, 3x/week) and Sex (female, male), and the within-subjects continuous variable Day. The random effects structure included an intercept (Subject) and slope (Day). For Experiment 2, predictors included the between-subjects categorical variables Group (daily intermittent, 3x/week) and Sex (female, male), the within-subjects categorical variable Cycle (Cycle 1, Cycle 2, Cycle 3), and the continuous variable Day. Planned comparisons were run for significant interactions including Group or Sex using Type III Sums of Squares. All data were analyzed using R (R 4.0.3). The highest order significant interactions (p < 0.05) including the variables Group and Sex were followed by planned comparisons using R’s emmeans or emtrends package. Results are visually presented as raw data mean ±SEM and the model fit mean ±SEM.

## Results

### Daily intermittent palatable food access leads to the most robust increase in binge-like eating

A Group × Sex × Day regression for PF intake showed a significant Group × Day interaction, (χ^2^(2) = 27.65, *p* < 0.01) and Group × Sex interaction, (χ^2^(2) = 9.29, *p* < 0.01). Inspection of the data leading to the Group x Day interaction indicated that mice in the daily intermittent group escalated their PF consumption across days (*t*(18.1) = 4.94, *p* < 0.01) compared to the continuous group (Figure 1A). There were also trends towards the 3x/week intermittent access group escalating their PF consumption compared to the continuous group (*t*(18.1) = 2.46, *p* = 0.06), and the daily intermittent group consuming more than the 3x/week group (*t*(17.5) = 2.47, *p* = 0.06). However, the daily intermittent group demonstrated the most robust increase in PF intake over time. Further examining potential sex differences in the behavioral response to PF access schedules, the Group × Sex interaction revealed that females in both the daily intermittent group (*t*(17.8) = 5.80, *p* < 0.01) and the 3x/week group (*t*(17.8) = 3.54, *p* < 0.01) consumed significantly more PF than the continuous group (Figure 1B). Additionally, females in the daily intermittent group trended towards consuming more PF than the 3x/week group (*t*(17.8) = 2.40, *p* = 0.07), consistent with the results in Figure 1A. In contrast, there were no differences in PF intake between groups in males (all *p* > 0.25; Figure 1B). Finally, there was a trend towards higher PF consumption in the daily intermittent group in females vs. males (*t*(17.8) = 1.98, *p* = 0.06) despite all PF intake being standardized by body weight prior to analysis. Together, these results support the presence of sex differences in this model, similar to BE in humans (Klump et al., 2017) and rats (Klump et al., 2013).

**Figure 1.**
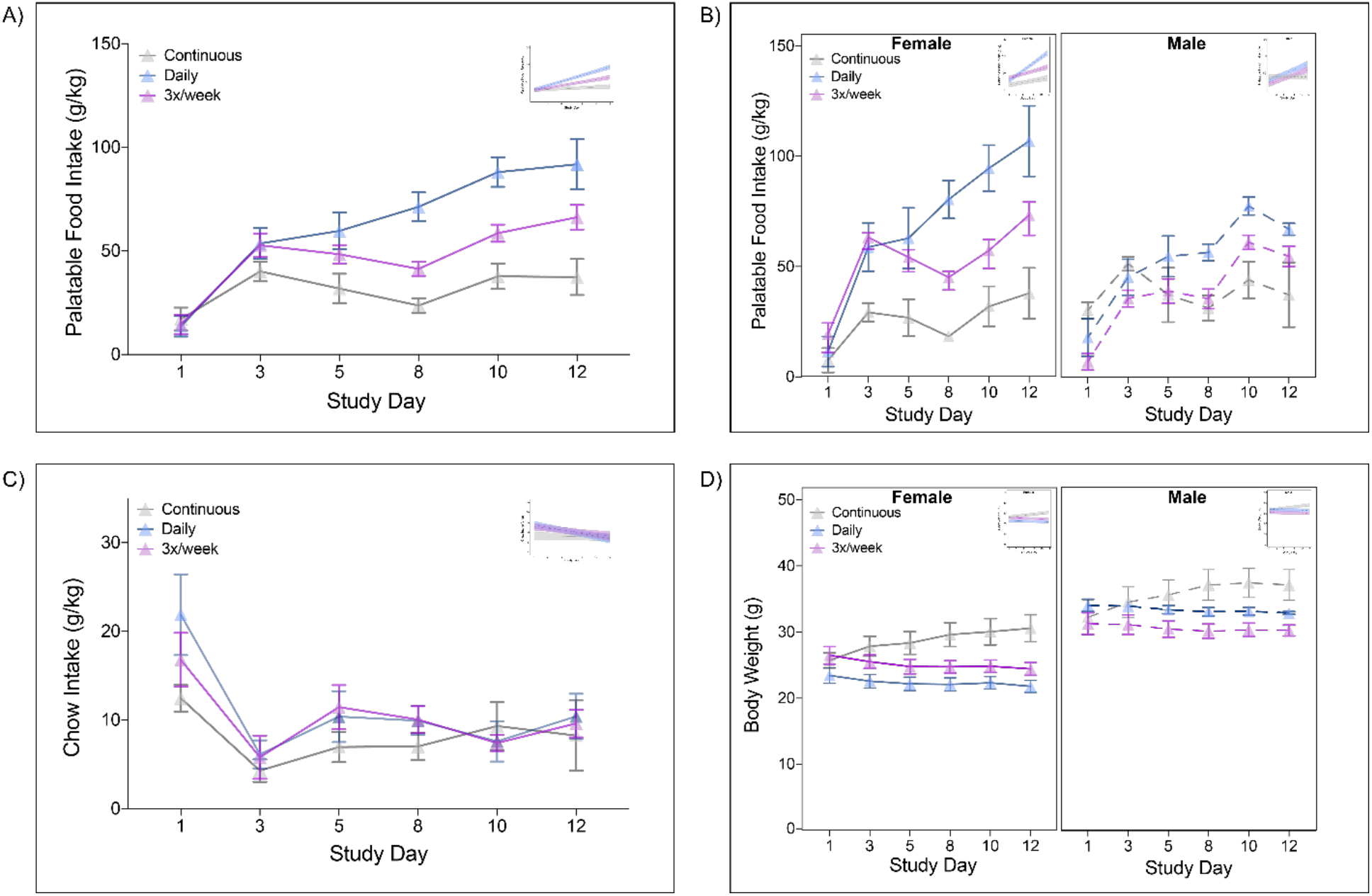
Early trajectories of palatable food intake, chow intake, and body weight across binge-like earing phenotypes. *Note*. All intake standardized by body weight (g/kg). PF &#x003D; palatable food. Sample sizes across groups: daily intermittent N &#x003D; 3 male, 5 female; 3x/week &#x003D; 3 male, 5 female; continuous &#x003D; 4 male, 4 female. Data shown is raw data. Inset in each figure represents model fit from regression analyses. A) The daily intermittent group (blue) significantly escalated PF consumption across days vs. the continuous group (gray) (*p* &#x003C; 0.01). B) females in both the daily intermittent group (blue) (*p* &#x003C; 0.01) and the 3x/week group (purple) (*p* &#x003C; 0.01) consumed significantly more PF than the continuous group. No differences in PF intake between males (all *p*&#x2019;s &#x003E; 0.25). C) Chow intake decreased across study days for all groups (*p* &#x003C; 0.01). D) Males weighed more than females (*p* &#x003C; 0.01), and both males and females in the continuous group showed increased weight over time (*p*&#x2019;s &#x003C; 0.01).

While results showed a sex difference in consumption of PF, there were no significant main effects or interactions for chow intake (including Group or Sex; all *p* > .1). However, there was a main effect of time, with chow intake decreasing across groups over study days (χ^2^(2) = 6.12, *p* < 0.01; Figure 1C). There was also a significant main effect of Sex for body weight showing that males weighed more than females as expected (χ^2^(1) = 43.04, *p* < 0.01; Figure 1D), and a significant Group × Day interaction (χ^2^(2) = 90.58, *p* < 0.01) such that body weight increased over time in the continuous group compared to the daily intermittent (*t*(18) = 8.04, *p* < 0.01) and 3x/week (*t*(18) = 8.29, *p* < 0.01) groups (Figure 1D).

### Patterns of binge-like eating persist across cycles of re-exposure to palatable food in mice with history of intermittent palatable food exposure

#### Baseline Phase

Results for PF intake during the Baseline Phase (i.e., first 12 days) of a new cohort largely replicated the findings reported in Figure 1A (Figure 2). There were significant Group × Day (χ^2^(1) = 24.55, *p* < 0.01) and Group × Sex (χ^2^(1) = 15.77, *p* < 0.01) interactions. The Group × Day interaction showed that the daily intermittent group escalated PF consumption across days more than the 3x/week group (*t*(11.4) = 4.61, *p* < 0.01). The Group × Sex interaction revealed that females consumed more PF than males in the daily intermittent group (*t*(11.6) = 5.21, *p* < 0.01), but there were no differences between males and females in the 3x/week group (*t*(11) = 0.10, *p* = 0.93) (Figure 2B, Baseline). Unlike the previous experiment where there was a decrease in chow intake over time, here there were no significant main effects or interactions of Group, Sex or Day in chow consumption (all *p* > 0.10).

**Figure 2.**
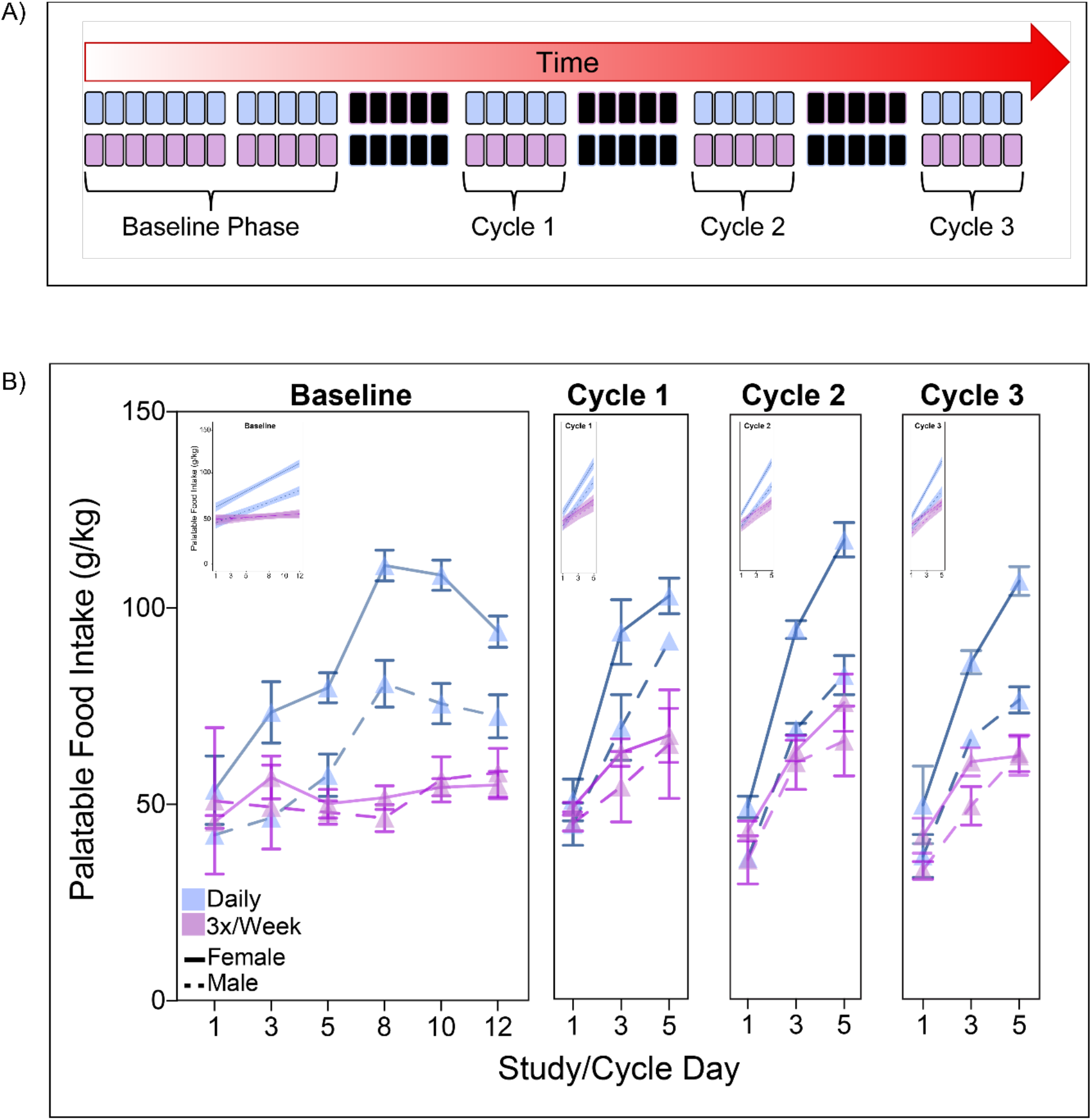
Patterns of palatable food intake across multiple incubation cycles in two binge-like intermittent schedules. *Note*. All intake standardized by body weight (g/kg). PF &#x003D; palatable food. Sample sizes across groups: daily intermittent N &#x003D; 3 male, 4 female; 3x/week &#x003D; 3 male, 5 female. Data shown is raw data. Inset in each figure represents model fit from regression analyses. A) Timeline for Baseline and Incubation cycles 1-3. B) During Baseline, the daily intermittent group escalated PF intake more than the 3x/week group (*p* &#x003C; .01), and females in the daily intermittent PF group consumed significantly more PF than males in the daily intermittent group (*p* &#x003C; .01). Patterns were similar across incubation cycles, with the daily intermittent group showing escalated PF intake more than the 3x/week group (*p* &#x003C; .01). Females also continued to consume the highest amount of PF (*p* &#x003C; .01).

In addition, there was a significant Group × Sex interaction for body weight, (χ^2^(1) = 4.30, *p* = 0.04). Planned comparisons showed that males weighed more than females in both the daily intermittent (*t*(11) = 4.73, *p* < 0.01), and the 3x/week groups (*t*(11) = 2.62, *p* = 0.02) as expected. While there was a significant Sex × Day interaction, (χ^2^(1) = 4.47, *p* = 0.03), planned comparisons found a nonsignificant trend towards males’ weight increasing more across days than females’ (*t*(11) = 2.06, *p* = 0.06).

#### Incubation Phase

There was a significant Group × Day interaction (χ^2^(1) = 27.85, *p* < 0.01) and Group x Sex interaction (χ^2^(1) = 4.58, *p* = 0.04) for PF intake across the incubation phase, showing that the daily intermittent group escalated their consumption across days more than the 3x/week group (*t*(10.9) = 4.88, *p* < 0.01), regardless of sex (Figure 2, Cycles 1-3). The Group x Sex interaction revealed that females consumed more PF than males within the daily intermittent group (*t*(11) = 4.19, *p* < 0.01), but not the 3x/week group (*t*(10.9) = 1.32, *p* = 0.21). Females in the daily intermittent group also consumed more PF than females in the 3x/week group (*t*(11.1) = 6.01, *p* < 0.01). This pattern of the daily intermittent group consuming more PF than the 3x/week group was also seen in the males, but it was only marginally significant (*t*(10.9) = 2.20, *p* = 0.0501). Interestingly, there was a significant Group × Sex × Cycle interaction (χ^2^(1) = 4.18, *p* = .04) found for chow intake. While planned comparisons showed no significant differences, results showed that the interaction was driven by males in the daily intermittent group which trended towards eating more chow during the second incubation cycle (*t*(10.84) = 1.92, *p* = 0.08).

Results showed a significant Group × Cycle interaction (χ^2^(2) = 16.89, *p* < 0.01), and Sex × Day interaction (χ^2^(2) = 19.45, *p* < 0.01) for body weight across cycles. The Sex × Day interaction showed that males’ body weights increased across days at a higher rate than females (*t*(11) = 4.44, *p* < 0.01). The Group × Cycle interaction showed that the groups did not differ within each cycle (all *p* > 0.10). However, weights increased across cycles within the 3x/week group (all *p* < 0.05). Body weight in the daily intermittent group increased between Cycle 1 and 2 (*t*(11) = 3.45, *p* = 0. 01), and Cycle 1 and 3 (*t*(11) = 4.96, *p* < 0.01), but not Cycle 2 and 3 (*t*(11) = 1.98, *p* = 0.16).

## Discussion

Pre-clinical approaches are uniquely suited to study the onset, development, and persistence of BE like behavior in a controlled setting. Here we examined three different PF access schedules in both male and female mice to identify a schedule that yields a robust and persistent pattern of binge-like eating. In the first 12 days of the BE paradigm, mice with daily intermittent access to PF escalated their intake more than mice in the 3x/week or continuous groups (Figure 1A, Figure 2B). This pattern was more pronounced in the female versus male mice (Figure 1B). Novel findings from the current study showed that not only did female mice in the daily intermittent group consume significantly more PF than female mice in the 3x/week group during the Baseline phase, this pattern re-emerged across multiple incubation cycles (Figure 2B, Cycles 1-3). This model may therefore serve as a critical tool to advance in our understanding of BE persistence in future studies. While, to our knowledge, this is the first investigation of pre-clinical BE persistence across cycles, models for substance use often use incubation/re-exposure models to examine persistence of drug use (Freeman et al., 2008; Gobin et al., 2019; Guillem et al., 2014), providing further support for the potential benefits of this paradigm in understanding BE persistence.

Our findings across experiments provide strong support for this model’s utility in quantifying BE onset and PF consumption trajectories, particularly in female mice. Results showed that mice with daily intermittent exposure to PF show the most robust increase in PF intake, extending previous work using daily intermittent PF access to elicit binge-like eating in rodents (e.g., Bake et al., 2013; Berner et al., 2008; Doucette et al., 2015). Additionally, female mice with daily intermittent PF access consumed significantly more PF than females with continuous access, consistent with findings in male rats where daily intermittent, rather than continuous, access resulted in higher PF intake (Furlong et al., 2014). Finally, daily intermittent access female mice consumed significantly more PF than daily intermittent access male mice, supporting previously identified sex differences in binge-like eating in rats (females > males) (Klump et al., 2013). Our results strongly align with the clinical presentation of BE–i.e. intermittent, rather than continuous, in nature, with episodes that are typically around two hours in duration (American Psychiatric Association, 2013). Additionally, there are sex differences observed in humans that engage in BE (females > males) (Klump et al., 2017; Striegel-Moore et al., 2009). In addition to modeling the clinical phenotype, this model to quantify early PF intake escalation may predict if an animal is at higher risk of persistent binge-like eating during the incubation phase. Identifying these early at-risk phenotypes may guide development of early interventions for BE, particularly in high-risk populations.

This work also advances our ability to model persistence of BE using pre-clinical approaches. Recent work found that during a one day re-exposure test after incubation of binge-like eating, mice consumed similar amounts or more PF compared to the last day of baseline tests, highlighting the potential to examine BE persistence (Sena et al., 2022). The current work expands on this finding by completing three cycles of incubation and re-exposure to PF across multiple days to examine not only the amount of PF consumed, but the patterns of reemergence and persistence in PF intake. This cyclic pattern mimics the remission/relapse observed in clinical BE, in which periods of abstinence from BE are followed by cycles of relapse after treatment (Fairburn et al., 1993). A similar pattern of abstinence/relapse has been used in studies investigating persistent substance use in pre-clinical models (Marchant et al., 2013). These models have provided important insight into environmental factors (Imperio et al., 2018) and neural mechanisms contributing to various types of relapse (Farrell et al., 2018), as well as pharmacological interventions for substance use (Baek et al., 2022). Our current model will lend itself to similar investigations of the mechanisms that contribute to BE persistence, improving our currently limited understanding of treatment targets for BE relapse and persistence.

In sum, these experiments identified a pre-clinical approach to quantify early escalation and persistence of BE that has strong translational relevance. For example, clinical research has shown that initial severity of BE Is associated with BE persistence at 1-year follow up (Stice et al., 2021). Our model could be used to identify the neural mechanisms contributing to the early escalation of severe binge-like eating in daily intermittent access female mice and identify neural factors contributing to BE persistence using techniques including *in vivo* calcium imaging and optogenetics. Ultimately, translation between pre-clinical and clinical research can lead to identification of unique mechanisms underlying BE persistence that can be effectively targeted in treatment.

## Acknowledgements

This work was supported by the BRAIN Initiative and the National Institutes of Health (F32MH118687, T32MH016804 to BH).

## References

Agh, T., Kovács, G., Pawaskar, M., Supina, D., Inotai, A., & Vokó, Z. (2015). Epidemiology, health-related quality of life and economic burden of binge eating disorder: a systematic literature review. Eating and Weight Disorders-Studies on Anorexia, Bulimia and Obesity, 20 (1), 1–12.

American Psychiatric Association. (2013). Diagnostic and Statistical Manual of Mental Disorders (5th ed.). American Psychiatric Association.

Baek, J. J., Kline, H., Deveau, C. M., & Yamamoto, B. K. (2022). Roflumilast treatment during forced abstinence reduces relapse to methamphetamine seeking and taking. Addiction Biology, 27(1), e13082.

Bake, T., Duncan, J. S., Morgan, D. G. A., & Mercer, J. G. (2013). Arcuate nucleus homeostatic systems are not altered immediately prior to the scheduled consumption of large, binge-type meals of palatable solid or liquid diet in rats and mice. Journal of Neuroendocrinology, 25(4), 357–371. https://doi.org/doi.org/10.1111/jne.12008

Berner, L. A., Avena, N. M., & Hoebel, B. G. (2008). Bingeing, self-restriction, and increased body weight in rats with limited access to a sweet-fat diet. Obesity, 16(9), 1998–2002. https://doi.org/10.1038/oby.2008.328

Boggiano, M. M., Artiga, A. I., Pritchett, C. E., Chandler-Laney, P. C., Smith, M. L., & Eldridge, A. J. (2007). High intake of palatable food predicts binge-eating independent of susceptibility to obesity: an animal model of lean vs obese binge-eating and obesity with and without binge-eating. International Journal of Obesity, 31(9), 1357–1367. https://doi.org/10.1038/sj.ijo.0803614

Doucette, W. T., Khokhar, J. Y., & Green, A. L. (2015). Nucleus accumbens deep brain stimulation in a rat model of binge eating. Translational Psychiatry, 5(12), 1–6. https://doi.org/10.1038/tp.2015.197

Fairburn, C. G., Peveler, R. C., Jones, R., Hope, R. A., & Doll, H. A. (1993). Predictors of 12-month outcome in bulimia nervosa and the influence of attitudes to shape and weight. Journal of Consulting and Clinical Psychology, 61(4), 696.

Farrell, M. R., Schoch, H., & Mahler, S. V. (2018). Modeling cocaine relapse in rodents: Behavioral considerations and circuit mechanisms. Progress in Neuro-Psychopharmacology and Biological Psychiatry, 87, 33–47.

Freeman, W. M., Patel, K. M., Brucklacher, R. M., Lull, M. E., Erwin, M., Morgan, D.,… Vrana, K. E. (2008). Persistent alterations in mesolimbic gene expression with abstinence from cocaine self-administration. Neuropsychopharmacology, 33(8), 1807–1817.

Furlong, T. M., Jayaweera, H. K., Balleine, B. W., & Corbit, L. H. (2014). Binge-like consumption of a palatable food accelerates habitual control of behavior and is dependent on activation of the dorsolateral striatum. Journal of Neuroscience, 34(14), 5012–5022. https://doi.org/10.1523/JNEUROSCI.3707-13.2014

Gobin, C., Shallcross, J., & Schwendt, M. (2019). Neurobiological substrates of persistent working memory deficits and cocaine-seeking in the prelimbic cortex of rats with a history of extended access to cocaine self-administration. Neurobiology of Learning and Memory, 161, 92–105.

Goldschmidt, A. B., Wall, M. M., Zhang, J., Loth, K. A., & Neumark-Sztainer, D. (2016). Overeating and binge eating in emerging adulthood: 10-year stability and risk factors. Developmental Psychology, 52(3), 475–483. https://doi.org/10.1037/dev0000086

Guillem, K., Ahmed, S. H., & Peoples, L. L. (2014). Escalation of cocaine intake and incubation of cocaine seeking are correlated with dissociable neuronal processes in different accumbens subregions. Biological Psychiatry, 76(1), 31–39.

Hildebrandt, B. A., & Ahmari, S. E. (2021). Breaking It Down: Investigation of Binge Eating Components in Animal Models to Enhance Translation. Frontiers in psychiatry, 1387. https://doi.org/10.3389/fpsyt.2021.728535

Hildebrandt, B. A., Klump, K. L., Racine, S. E., & Sisk, C. L. (2014). Differential strain vulnerability to binge eating behaviors in rats. Physiology & Behavior, 127, 81–86. https://doi.org/10.1016/j.physbeh.2014.01.012

Hildebrandt, B. A., Sinclair, E. B., Sisk, C. L., & Klump, K. L. (2018). Exploring reward system responsivity in the nucleus accumbens across chronicity of binge eating female rats. International Journal of Eating Disorders, 51, 989–993. https://doi.org/10.1002/eat.22895

Hudson, J. I., McElroy, S. L., Ferreira-Cornwell, M. C., Radewonuk, J., & Gasior, M. (2017). Efficacy of lisdexamfetamine in adults with moderate to severe binge-eating disorder: a randomized clinical trial. JAMA psychiatry, 74(9), 903–910.

Imperio, C. G., McFalls, A. J., Hadad, N., Blanco-Berdugo, L., Masser, D. R., Colechio, E. M.,… Vrana, K. E. (2018). Exposure to environmental enrichment attenuates addiction-like behavior and alters molecular effects of heroin self-administration in rats. Neuropharmacology, 139, 26–40.

Keski-Rahkonen, A. (2021). Epidemiology of binge eating disorder: prevalence, course, comorbidity, and risk factors. Current opinion in psychiatry, 34 (6), 525–531.

Kjeldbjerg, M. L., & Clausen, L. (2021). Prevalence of binge-eating disorder among children and adolescents: a systematic review and meta-analysis. European Child & Adolescent Psychiatry, 1–26.

Klump, K. L., Culbert, K. M., & Sisk, C. L. (2017). Sex differences in binge eating: Gonadal hormone effects across development. Annual Review of Clinical Psychology, 13, 183–207. http://www.annualreviews.org/doi/full/10.1146/annurev-clinpsy-032816-045309?url_ver=Z39.88-2003&rfr_id=ori%3Arid%3Acrossref.org&rfr_dat=cr_pub%3Dpubmed

Klump, K. L., Racine, S. E., Hildebrandt, B., & Sisk, C. L. (2013). Sex differences in binge eating patterns in male and female adult rats. International Journal of Eating Disorders, 46(7), 729–736. https://doi.org/10.1002/eat.22139

Marchant, N. J., Li, X., & Shaham, Y. (2013). Recent developments in animal models of drug relapse. Current Opinion in Neurobiology, 23(4), 675–683.

Pope, H. G., Lalonde, J. K., Pindyck, L. J., Walsh, B. T., Bulik, C. M., Crow, S. J, Hudson, J. I. (2006). Binge eating disorder: a stable syndrome. American Journal of Psychiatry, 163(12), 2181–2183.

Safer, D. L., Lively, T. J., Telch, C. F., & Agras, W. S. (2002). Predictors of relapse following successful dialectical behavior therapy for binge eating disorder. International Journal of Eating Disorders, 32(2), 155–163.

Sena, K. D., Beierle, J. A., Richardson, K. T., Kantak, K. M., & Bryant, C. D. (2022). Assessment of Binge-Like Eating of Unsweetened vs. Sweetened Chow Pellets in BALB/c Substrains. Frontiers in Behavioral Neuroscience, 16, 944890.

Solmi, M., Radua, J., Olivola, M., Croce, E., Soardo, L., Salazar de Pablo, G.,… Kim, J. H. (2022). Age at onset of mental disorders worldwide: large-scale meta-analysis of 192 epidemiological studies. Molecular Psychiatry, 27(1), 281–295.

Stice, E., Bohon, C., Gau, J. M., & Rohde, R. (2021). Factors that predict persistence versus non-persistence of eating disorder Symptoms: A prospective study of high-risk young women. Behaviour Research and Therapy, 144, 103932.

Striegel-Moore, R. H., Rosselli, F., Perrin, N., DeBar, L., Wilson, G. T., May, A., & Kraemer, H. C. (2009). Gender difference in the prevalence of eating disorder symptoms. International Journal of Eating Disorders, 42(5), 471–474.

Vall, E., & Wade, T. D. (2015). Predictors of treatment outcome in individuals with eating disorders: A systematic review and meta-analysis. International Journal of Eating Disorders, 48(7), 946–971.

